# Genomic adaptation of giant viruses in polar oceans

**DOI:** 10.1101/2023.02.09.527846

**Authors:** Lingjie Meng, Tom O. Delmont, Morgan Gaïa, Eric Pelletier, Antonio Fernàndez-Guerra, Samuel Chaffron, Russell Y. Neches, Junyi Wu, Hiroto Kaneko, Hisashi Endo, Hiroyuki Ogata

## Abstract

Despite being perennially frigid, polar oceans form an ecosystem hosting high and unique biodiversity. Various organisms show different adaptative strategies in this habitat, but how viruses adapt to this environment is largely unknown. Viruses of phyla *Nucleocytoviricota* and *Mirusviricota* are groups of eukaryote-infecting large and giant DNA viruses with genomes encoding a variety of functions. Here, by leveraging the Global Ocean Eukaryotic Viral database, we investigate the biogeography and functional repertoire of these viruses at a global scale. We first confirm the existence of an ecological barrier that clearly separates polar and nonpolar viral communities, and demonstrate that temperature drives dramatic changes in the virus–host network at the polar/nonpolar boundary. Ancestral niche reconstruction suggests that adaptation of these viruses to polar conditions has occurred repeatedly over the course of evolution, with polar-adapted viruses in the modern ocean being scattered across their phylogeny. Numerous viral genes are specifically associated with polar adaptation, although most of their homologues are not identified as polar-adaptive genes in eukaryotes. These results suggest that giant viruses adapt to cold environments by changing their functional repertoire, and this viral evolutionary strategy is independent of the polar adaptation of their hosts.

## Main

Polar regions are recognized as among the coldest environments on Earth, with strong seasonal variations in light cycles. Nevertheless, these regions could nourish a diverse range of creatures, from microscopic organisms to large animals, thanks to the primary production by phytoplankton. Organisms adapted to polar environments exhibit distinctive physiological or morphological characteristics, which augment their fitness in these extreme but lush environments. For example, polar bears show characteristic morphological traits whose underlying genetic variations occurred in their ancestral gene pools^1^. Both Arctic and Antarctic fishes encode antifreeze proteins that allow them to maintain physiological activity in cold waters^2,3^, while some psychrophilic bacteria produce oxygen-scavenging enzymes or modify their membrane fatty acid composition^4,5^.

How do viruses adapt to polar environments? In the ocean, viruses are the most abundant biological entities^6^ and play important roles in the regulation of microbial host communities, carbon and nutrient cycling, and horizontal gene transfer among organisms^7–10^. Recent metagenomic studies have revealed that both Arctic^11,12^ and Antarctic^13,14^ environments harbour diverse viruses, with an elevated diversity of prokaryotic dsDNA viruses in the Arctic Ocean^11^. A large proportion of Arctic-specific genes from these viruses were suggested under positive selection based on their mutation patterns. This implied a role for gene repertoire in viral adaptation, although most of those genes were of unknown function. It is also known that phylogenetically closely related viruses can display different responses in their infection dynamics to varying temperature^15,16^, suggesting that virus–host systems adapt to thermal changes. Another study showed that a prokaryotic virus reduced its genome in response to decreased culture temperature^17^. These studies imply possible adaptive mechanisms of viruses to low temperature or polar ecosystems. However, our knowledge on such viral adaptations is still limited.

In our previous study, we revealed a remarkable shift in the community composition of eukaryotic dsDNA viruses from nonpolar to polar biomes^18^. These viruses, classified in phylum *Nucleocytoviricota* (“giant viruses”), are known for their large genomes encoding hundreds to thousands of genes^19,20^. These viruses are ancient^21^, diverse^22,23^, abundant^24,25^, and active^26^ in the ocean. Despite the existence of a clear polar/nonpolar barrier for these viruses, the underlying genomic and ecological mechanisms are unknown. How frequently these viruses have crossed this polar barrier over evolutionary time also remains unclear. As the genomes of *Nucleocytoviricota* dynamically evolve by losing and gaining different functions^19,27^, we hypothesized that adaptation to polar environments impacts their gene repertoire.

In this study, we investigated genomes of eukaryotic large DNA viruses to characterize their genome-level adaptations to polar environments. We leveraged recently reconstructed viral and eukaryotic environmental genomes from the multidisciplinary *Tara* Oceans international research project^28,29^. The viral genomic data include environmental genomes of viruses from phylum *Nucleocytoviricota* and a recently discovered phylum, *Mirusviricota*^28^.

We first assess the existence of a polar barrier for giant viruses by analysing viral community composition and by computing robust temperature optima for viruses and their predicted hosts. We then perform ancestral state reconstruction for polar and nonpolar niches along the phylogenomic tree of these viruses to quantitatively estimate the adaptive evolutionary events. Finally, we delineate the functions that are specific to “polar” viruses and present evidence that viral genomic adaptation to low temperature is independent from the adaptation of their hosts.

## Results and Discussion

### Polar barrier for giant viruses

We investigated the biogeography of giant virus genomes from the Global Ocean Eukaryotic Viral (GOEV) database^28^. Their abundance profiles across *Tara* Oceans samples from different size fractions (Supplementary Fig. 1a,b; Supplementary Table 1) revealed 1,380 viral genomes that showed signals (>25% of the genome length was mapped by reads, see methods in our previous paper^28^) in at least one sample out of 928 samples (The details of biogeography were in the supplementary text; Supplementary Fig. 1-3). The presence/absence distribution of viral genomes across biomes revealed a large number of genomes specific to the Polar biome. Out of 569 genomes detected in polar regions, 262 (46.05%) were exclusive to the Polar biome (Supplementary Fig. 4a). Accordingly, biome-based classification of viral communities (i.e., Polar, Coastal, Trades, and Westerlies) had significant explanatory power for community variation (Supplementary Fig. 4b,c; ANOSIM, *P* < 0.01). The *R* value increased from 0.4021 to 0.6141 after merging three nonpolar biomes, demonstrating the existence of a clear polar barrier for giant virus communities. The viral communities of Arctic regions were also characterized by their relatively high abundances showing peaks in cumulative coverage plots for different size fractions (Supplementary Fig. 1b). The major groups of viruses in this area were *Algavirales*, followed by *Imitervirales* as in other areas of the ocean (Supplementary Fig. 2c).

We inferred a virus–plankton network through co-occurrence analysis to further characterize the polar barrier in the context of virus-host interaction. In this analysis, we combined our virus genome data with previously reconstructed marine eukaryotic genome data^29^. In total, 2,135 virus–eukaryote associations (edges) were inferred in the network, with the majority (91.94%) of them being positive associations (Fig. 1a; Supplementary Table 3). Virus–eukaryote pairs with strong associations (edge weight ≥ 0.4) showed significantly higher protein similarities between their genomes than those without strong associations (no edges or edges with weight <0.4) (Wilcoxon rank-sum test, *P* = 1×10^−13^) (Fig. 1b). Such an increase of sequence similarity can be due to horizontal gene transfers between viruses and hosts^30,31^, supporting the prediction of true virus–host relationships in the reconstructed network. A previous study revealed that the structure of the network for marine eukaryotes and prokaryotes correlates with the temperature optima of species^32^. By estimating robust temperature optima for individual viruses and eukaryotes^33^, we identified a strong correlation between the temperature optima and the structure of the virus–eukaryote network (Fig. 1a). A dramatic structural change in the network at the temperature-dependent polar/nonpolar boundary is the source of the uniqueness of polar viral communities.

**Fig. 1.**
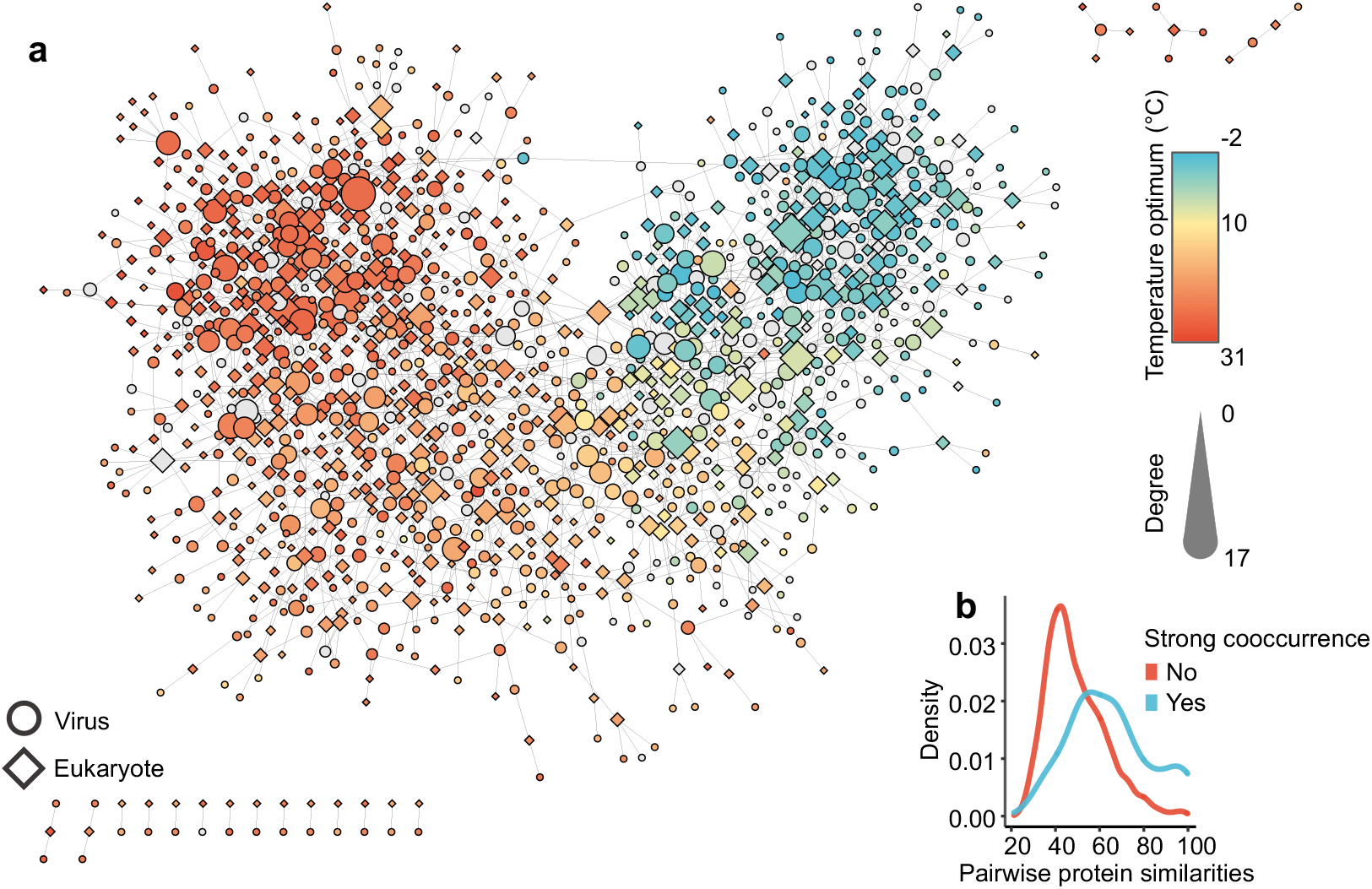
Virus–plankton interaction network. **a,** Five individual networks inferred using input matrices for the relative frequencies of eukaryotes (five size fractions) and giant viruses (Pico-size fraction). The best positive or negative association (i.e., the edges with the highest absolute weights between two genomes) were selected to build the integrated network. Node colour represents the temperature optima of each genome for viruses and eukaryotes. A total of 1,347 nodes (567 eukaryotes and 780 viruses) are in the network. Of these nodes, 1,191 nodes (554 eukaryotes and 637 viruses) are coloured according to their temperature optima. **b,** The distribution of pairwise sequence similarity of proteins (one protein from the eukaryotic genome and one from the viral genome). Blue line indicates the distribution for pairs with a strong virus–eukaryote association in the network (edge weight of ≥ 0.4), while the red line is for pairs lacking a strong association. The two distributions are significantly different (*P* = 1×10^−13^, Wilcoxon signed-rank test).

Latitudinal diversity gradients are characterized by relatively low polar and high temperate biodiversity^34^ and are widespread across all ranges of marine microorganisms^35^. Previous studies revealed a similar latitudinal diversity gradient for giant viruses^35^, but not for prokaryotic dsDNA viruses^11,35^ and RNA viruses^36^. In this study, various diversity gradient patterns were observed among viruses of different size fractions and main taxonomic groups (Supplementary Fig. 1d; Supplementary Fig. 5). The reasons underlying the Arctic diversity hotspots for some viruses (e.g., viruses in large size fractions and mirusviruses) may reflect their host ranges as previously suggested^35^. Notably, eukaryotic nodes (i.e., potential hosts) associated with viruses showed a pattern distinct from the general diversity gradient trend with increasing diversity towards high latitudinal regions (Supplementary Fig. 6).

### Potential hosts for polar viruses

A phylogeny-informed filtering method, Taxon Interaction Mapper (TIM)^37,38^, was applied to the edges of the network to assign predicted host taxa to viral clades. This method assigned predicted host taxa (five taxa in total) to 34 viral clades (Supplementary Fig. 7a). These predictions are summarized in Supplementary Fig. 7b and included known virus–host relationships: *Mesomimiviridae* (from *Imitervirales*) and Phaeocytales^39–41^; *Mesomimiviridae* and Pelagomonadales^42,43^; and *Prasinovirus* (from *Algavirales*) and Mamiellales^44,45^.

Recent discoveries of giant endogenous viral elements (GEVEs) that are widespread across different eukaryotes demonstrated the impacts of giant viruses’ infection on host genome evolution^46–49^. We systematically analysed insertions of genomes of giant viruses and their satellite viruses (i.e., virophages) in marine eukaryotic genomes^29^. Among the five eukaryotic taxa predicted to contain viral hosts, the diatom order Chaetocerotales showed the largest number of insertion signals of both giant viruses and virophages (Supplementary Fig. 7b), suggesting infection of dsDNA viruses in these diatoms. Because only ssDNA and ssRNA viruses have been reported to infect species of diatoms^50^, we further analysed draft genomes of two isolated *Chaetoceros* species^51,52^, revealing three putative GEVEs in *C. tenuissimus* and one GEVE in *C. muelleri*. Two viral DNA polymerase genes detected in the *Chaetoceros* genomes were phylogenetically placed close to *Asfuvirales* and *Imitervirales* clades (Supplementary Fig. 7c), corroborating the virus–host relationships of *Imitervirales*/Chaetocerotales and *Asfuvirales*/Chaetocerotales predicted by our phylogeny-informed co-occurrence method. Because chaetocerotalid diatoms are abundant and diverse in both the Arctic and Southern Oceans^53,54^, this unidentified virus–host relationship may account for the diversity of giant viruses in high-latitude regions.

### Recurrent polar adaptations throughout viral evolution

To investigate viral adaptation across the polar barrier, we assigned ecological niche categories, either “Polar” or “Nonpolar”, to individual viral genomes. Of 1,380 viral genomes, 450 genomes were classified as Polar, while 818 genomes were classified as Nonpolar (Fig. 2a,b). 111 genomes were labelled “Unknown” because of their ambiguous distribution patterns. This ecological niche assignment was consistent with the robust temperature and latitude optima (Supplementary Fig. 8a). We then investigated the niche category assignment in the phylogenomic tree of viruses and found numerous clades of Polar viruses scattered across the tree (Fig. 3a). One Polar clade included an Arctic-original metagenome-assembled genome (MAG) and organic lake phycodnaviruses derived from an Antarctic organic lake^14^ (Supplementary Fig. 8b). *Emiliania huxleyi* viruses, known to occur at high latitudes, were also assigned to Polar clades. All six genomes of *Proculviricetes*^28^, a recently discovered class-level group recovered exclusively from the Arctic and Southern Oceans, were classified as Polar viruses. These examples corroborate the reliability of the ecological niche assignment using global-scale abundance profiles.

**Fig. 2.**
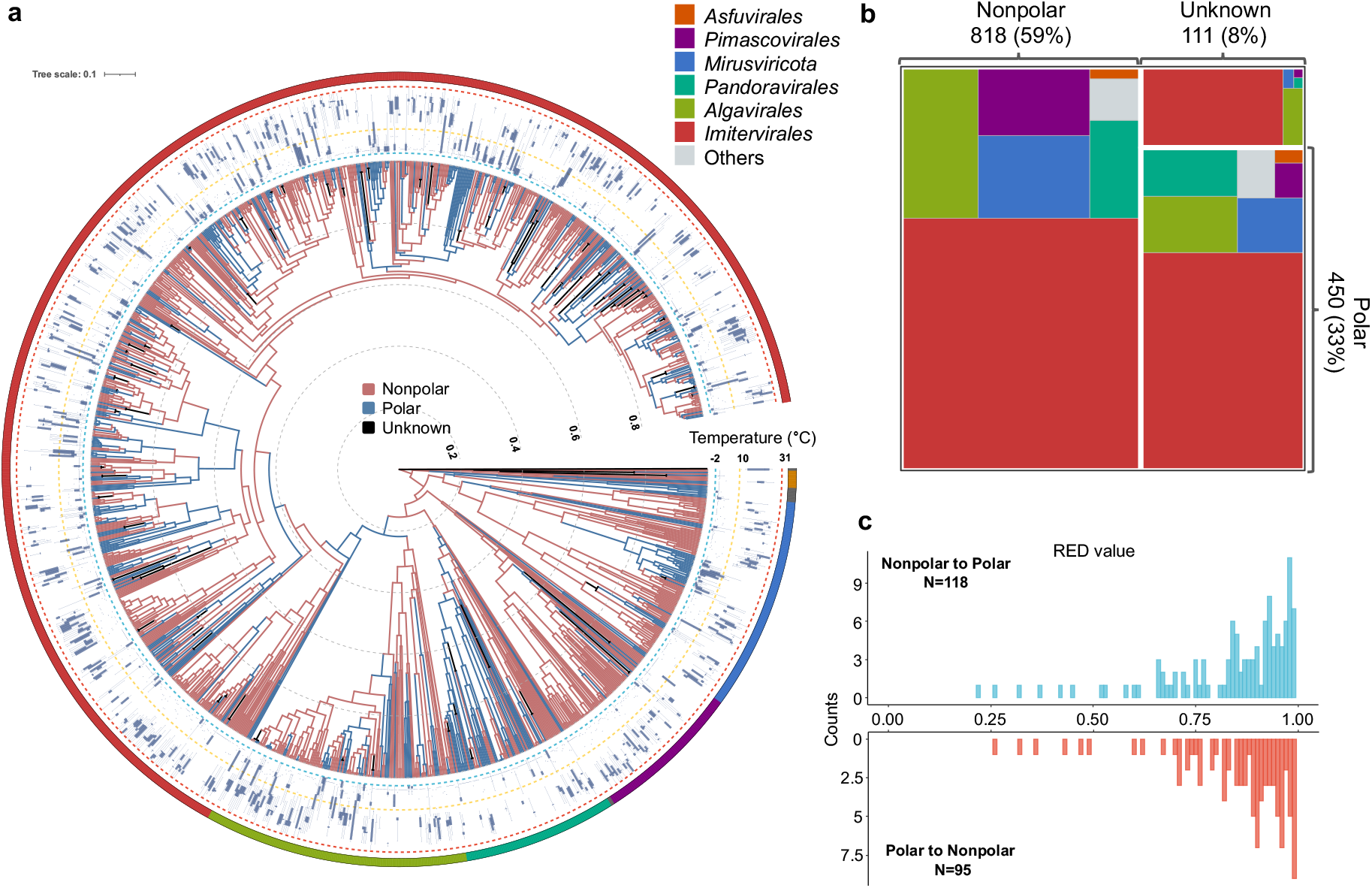
Inferred ancestral polar and non-polar niches for viruses. **a,** Ancestral “Polar” and “Nonpolar” states were estimated using the phylogenetic tree based on a one-parameter equal rates model. The outermost layer shows the taxonomy of six main groups. The boxplots in the second layer show the temperature optima of the viral genomes. Only polar and nonpolar genomes were included in the tree. **b,** The treemap diagram shows the number of viruses assigned to Polar, Nonpolar or “Unknown” biomes. Colours indicate the main taxonomic groups. **c,** Histograms of Relative Evolutionary Divergence (RED) values for the nodes at which “polar” or “nonpolar” adaptation events were inferred.

**Fig. 3.**
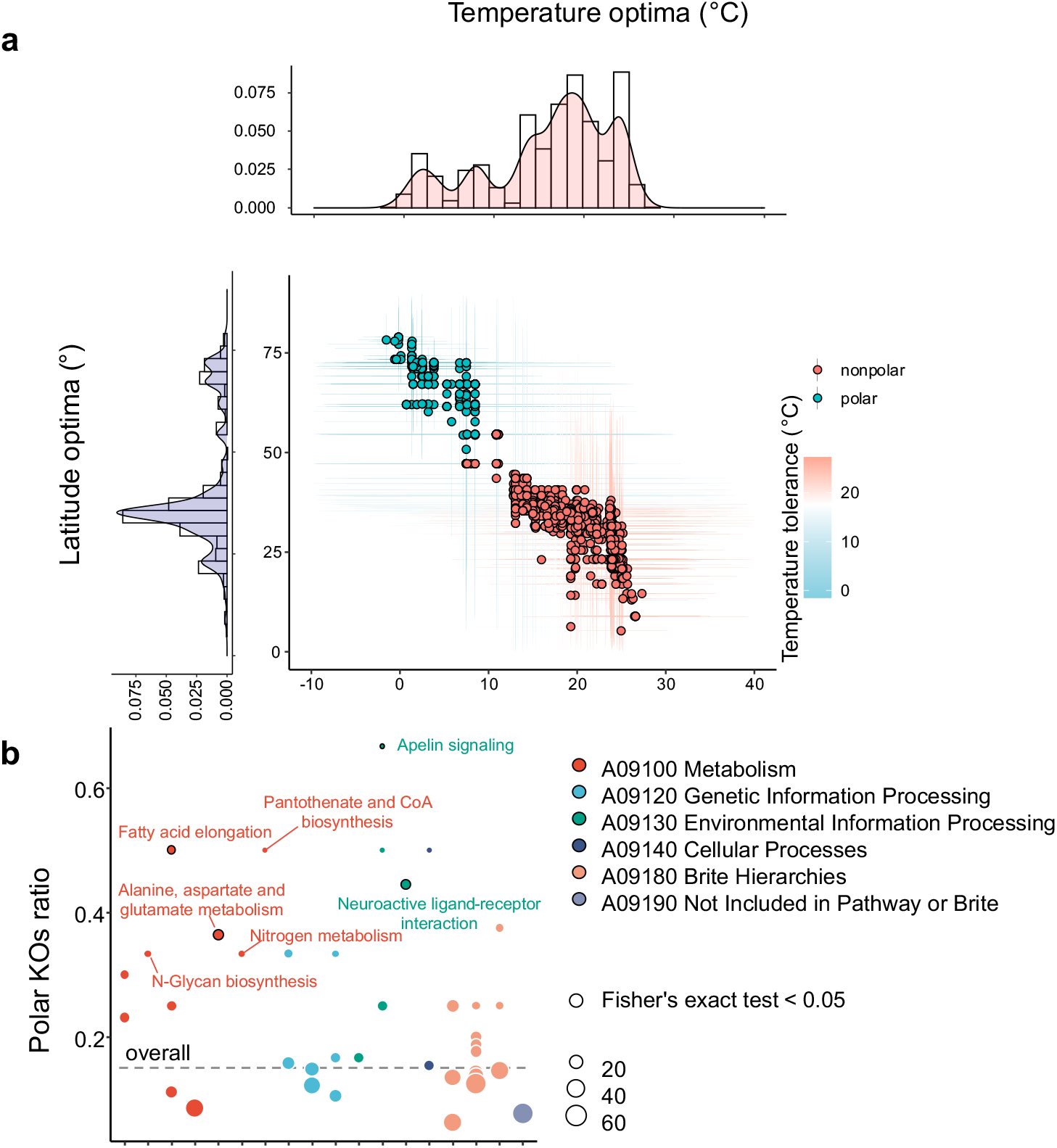
Ecological niche of KEGG Orthologs (KOs) and polar-enriched pathways. **a,** Distribution of the temperature optima and latitude optima for KEGG Orthologs (KOs) found in viral genomes. Colours of dots represent the Polar or Nonpolar niche for each KO. Bars indicate the tolerance ranges of temperature (horizontal) and latitude (vertical). Histograms show the distributions of temperature and latitude optima. **b,** Ratio of Polar KOs in each pathway. Black-framed circles correspond to pathways in which Polar KOs were significantly enriched (*P* < 0.05, Fisher’s exact test). The overall ratio of Polar KOs to all KOs is indicated by a dotted line.

We then performed Polar/Nonpolar state reconstruction for ancestral nodes in the tree using a maximum likelihood approach (see Methods). As a result, 118 Nonpolar-to-Polar and 95 Polar-to-Nonpolar niche adaptations were inferred along the branches of the tree (Fig. 2a). These adaptations thus occurred recurrently throughout the evolution of these viruses starting from the root of the tree, which was inferred as Nonpolar. Yet, our data could not exclude the possibility of a polar-origin scenario due to the difficulty in determining the root of the tree of giant viruses. The divergence of these viruses is estimated to predate the divergence of eukaryotes^21,23^. Most of the reconstructed niche adaptations occurred relatively recently after the formation of genera, but some adaptations were inferred to have occurred during the early stage of evolution, corresponding to order-level divergence (Fig. 2c).

### Polar-specific viral functions and their phylogenetic distributions

Genomic adaptation (i.e., adaptation by alteration of gene repertoire) to polar regions was investigated based on functions encoded in the viral genomes. We first annotated genes in the viral genomes with the KEGG Orthologs (KOs). For KOs (n = 1591) that were observed in more than four genomes, we calculated robust temperature and latitude optima (Supplementary Table 4). The temperature optima ranged from −1.54 °C to 27.31 °C, and the latitude optima from 5.25 ° to 78.96 °. The distribution of these values revealed two major groups of KOs: one distributed in high-latitude/low-temperature regions (n = 314, 19.74%) and another in lower-latitude/higher-temperature regions (n = 1,277, 80.26%) (Fig. 3a). The 314 Polar-specific genes had temperature optima below 10 °C and latitude optima above 50 °. The temperature and latitude optima for conserved core genes of giant viruses were found in the second group, being distributed at around 13–14 °C and 37–40 °, respectively.

We then calculated the phylogenetic diversity of individual KOs using the viral phylogenomic tree as a reference to assess the breadth of their phylogenetic distribution (Supplementary Fig. 9a). Overall, Polar-specific KOs showed a relatively low phylogenetic diversity (median = 6.94) compared with other KOs (median = 9.67) (Wilcoxon rank-sum test, *P* < 0.01), indicating relatively narrow phylogenetic distributions of the Polar-specific KOs. To further characterize the phylogenetic distributions of the 314 Polar-specific KOs, we examined the strength of phylogenetic signals in their distribution using a model comparison approach (see Methods). This analysis revealed that the reference phylogenomic tree has insufficient explanatory power for the phylogenetic distribution of 193 Polar-specific KOs (61%) out of the 314 KOs (chi-squared test, *P* < 0.05). It is thus inferred that additional factors rather than speciation history impacted the phylogenetic distribution of these KOs; environmental conditions or associated host distributions could be such factors.

### Polar-specific viral functions and metabolic pathways

The proportion of polar-specific KOs (among all genes with KO annotations in a viral genome) was significantly higher in Polar genomes (15.84% on average) compared to Nonpolar (6.95%) and Unknown (7.93%) genomes (Supplementary Fig. 9b; Kruskal-Wallis test, *P* < 0.01). Among Polar-specific KOs, ceramide glucosyltransferase (K00720) and dihydrofolate reductase (K18589) were exclusively distributed in polar genomes. Ceramide glucosyltransferase catalyzes sphingolipid glycosylation, indicating the biosynthesis of viral sphingolipids may improve the fitness of polar viruses^55^. Dihydrofolate reductase could provide dTMP pools for low GC content viruses, and a possible role of this function is to facilitate the replication of viruses in the persistent infections^56^. Additionally, nitrate transporter (K02575) had a high ratio of polar to nonpolar phylogenetic diversity (ratio = 7.96), thus showing a comparatively wide phylogenetic distribution in Polar genomes. The nitrate transporter pathway has a role in assimilating extracellular nitrate/nitrite, implying a potential role for Polar viruses to reprogram host metabolism to fit the nitrate-deficient polar oceans^57^. Some metabolic functions, including CoA biosynthesis (4’-phosphopantetheinyl transferase) and secondary metabolite biosynthesis (hydroxymandelonitrile lyase and 2-polyprenyl-6-hydroxyphenyl methylase), also showed a high phylogenetic diversity for Polar genomes.

At the pathway level, we found that six pathways were significantly enriched in Polar KOs (Fig. 3b; Fisher’s exact test, *P* < 0.05). Biosynthesis of unsaturated fatty acids, was found to be the most significantly enriched with polar KOs. A high proportion of unsaturated fatty acids is known as an adaptive trait for bacteria inhabiting low temperature and high pressure environment^58^. Giant viruses isolated from high latitude areas are known to encode enzymes for the biosynthesis of unsaturated fatty acid^41^ and may rewire the host physiology of fatty acid^55^. The N-glycan biosynthesis pathway also had a relatively high ratio of Polar-specific KOs. N-glycan influences the virus replication cycle, including virus recognition and virus release^59^. Neuroactive ligand-receptor interaction was the pathway most significantly enriched with polar specific KOs, implying the ability of polar viruses to regulate signal transduction. Collectively, these results underscore the importance of membrane-related pathways, including unsaturated fatty acid and specific membrane-related functions, in polar virus–host interactions.

### Other potential polar adapted functions

In addition to the above statistical analyses based on the temperature and latitude optima, we performed an enrichment analysis of KOs by examining their presence in Polar and Nonpolar genomes at different evolutionary scales to capture a variety of situations in the phylogenetic distributions of KOs. Specifically, this analysis was performed at four different lineage levels (i.e., root, main group, family, and genus). The analysis revealed 265 functions that were significantly enriched in Polar genomes inside at least one lineage (Fisher’s exact test, *P* < 0.05; Supplementary Table 4). These KOs enriched in Polar viral genomes showed lower temperature optima than other KOs (Supplementary Fig. 9c; Wilcoxon rank-sum test, *P* < 0. 01). For a finer-grained observation, we focused on one *Mesomimiviridae* clade, containing a similar number of Polar (n = 32) and Nonpolar (n = 40) genomes scattered in a subtree of the phylogenomic tree. In this example, four functions were found in more than five genomes from different Polar clades (Fig. 4a). Three of them (K01627, K00979, K06041) co-occurred in the same genomes and formed a near-complete CMP-KDO biosynthesis module in the lipopolysaccharide biosynthesis pathway (Fig. 4b). Lipopolysaccharides are the main component of the Gram-negative bacterial outer membrane, and enzymes of CMP-KDO biosynthesis were found in the genome of Cafeteria roenbergensis virus^60^. This result suggests that the genomes in the examined Polar clade have adapted to the polar environment by coating virions with bacteria-like glycoconjugate to enhance their interactions with Polar hosts.

**Fig. 4.**
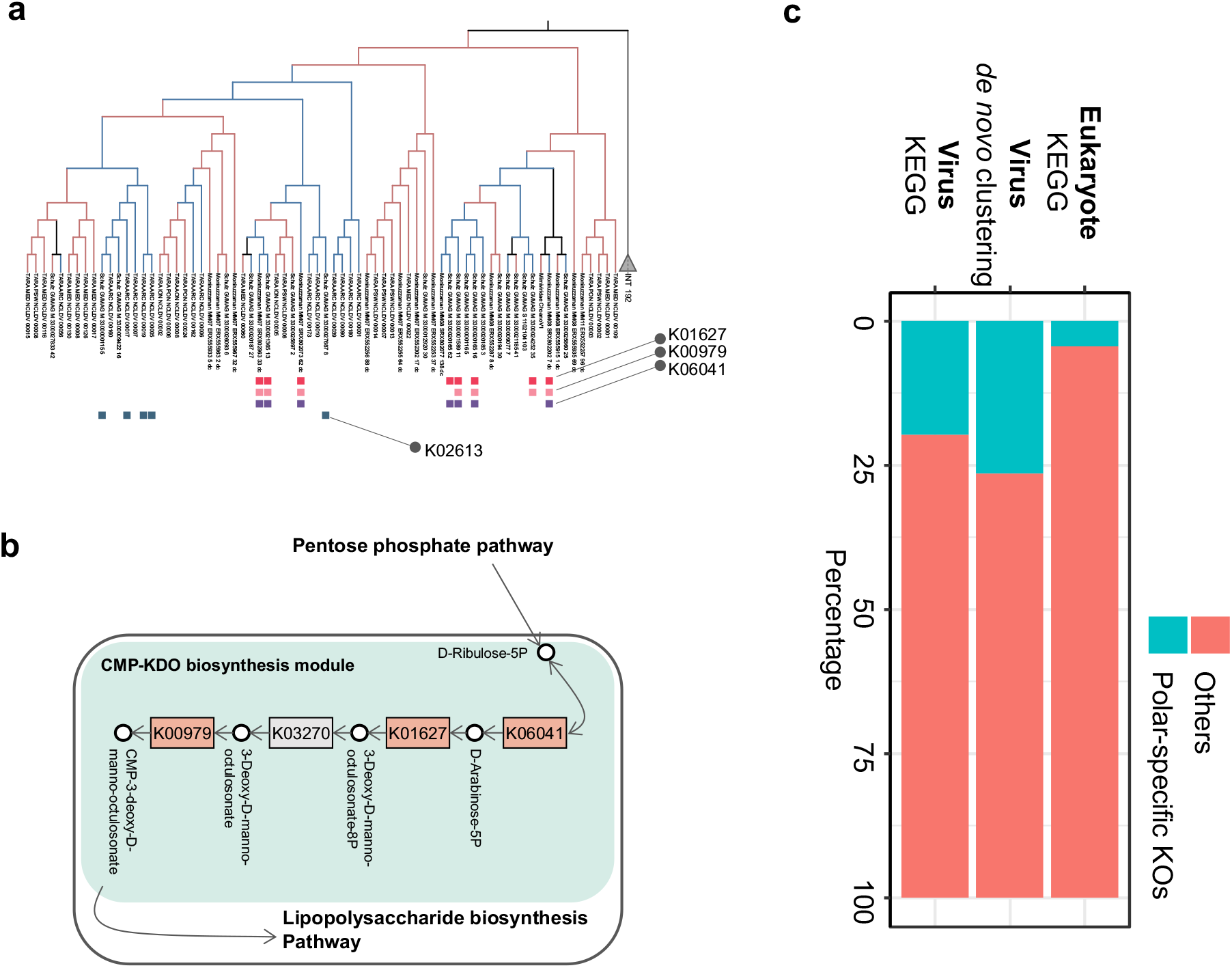
Independent genomic adaptation of giant viruses. 244 functions (KOs) were enriched at individual lineages. One example was given in **a,** Four KOs that were present exclusively in more than five Polar genomes in a selected *Mesomimiviridae* clade. Three of them (K01627, K00979, K06041) were encoded in the same genomes and formed a near-complete CMP–KDO biosynthesis module shown in **b,** Schematic of the three Polar enzymatic steps in the CMP–KDO biosynthesis module. **c,** Proportion of Polar and Nonpolar specific functions (KOs and GCCs) in viruses and eukaryotes.

The KO system can annotate only functionally known genes, and therefore we calculated robust temperature and latitude optima for gene cluster communities, *de novo* clusters of viral genes^28^. The result indicated a slightly higher proportion of Polar-specific gene clusters (26.43%) than obtained by KO annotations (19.74%) (Fig. 4c; Supplementary Fig. 10a), indicating the presence of genes of unknown function that show Polar-specific distributions. We also found that Polar genomes have a slightly but significantly higher proportion of Alanine-rich low-complexity regions than Nonpolar and Unknown genomes (Supplementary Fig. 9d; Dunn’s test, *P* < 0.05, following a significant Kruskal-Wallis test, *P* = 0.0002). These low-complexity sequences potentially have an anti-freeze function, as alanine-rich helical structure is one of the significant characteristics of type I antifreeze proteins for ice growth inhibition^61^. Additionally, the proportion of Polar viral genomes that encoded antifreeze protein homologs (n = 7, 1.6%) was higher than the genomes of other groups (n = 6, 0.65%), although the difference was not statistically significant (*P* > 0.05).

### Polar-specific functions in microbial-eukaryotes

Finally, to examine whether genomic adaptation occurs in eukaryotic plankton in polar regions and to test if the adaptation is related to the one in viruses, we calculated the temperature and latitude optima for KOs (n = 11,988) assigned to genes in eukaryotic genomes. A similar pattern of Polar and Nonpolar KO groups was identified, although the proportion of the Polar KO group (n = 523, 4.36%) was much smaller than that for viruses (19.74%) (Fig. 4c; Supplementary Fig. 10b). Interestingly, of the 523 KOs in the eukaryotic Polar group, only four were found in the viral Polar group. These were PPM family protein phosphatase, L-galactose dehydrogenase, transcription factor S, and ATP-dependent DNA helicase DinG. This result indicates that most Polar viral functions do not exhibit the same temperature/latitude optima seen in eukaryotic genomes. The result further suggests that virus–host horizontal gene transfer is not the primary driver of viral polar adaptation, and that genomic adaptations are uncoupled between viruses and eukaryotes.

## Conclusions

Functional repertoire is considered an important trait for the adaptation of organisms. Previous discoveries of functionally related genes in viruses^55,62^ indicated that functional repertoire could also be important for adaptive evolution of viruses. However, this has rarely been addressed for large and giant DNA viruses at a wide geographic scale as compared with cellular organisms. Thanks to the recent progress in metagenomics, we investigated the links between the biogeography, host types, and gene repertoire of viruses infecting marine eukaryotes. We confirmed the existence of a strong polar/nonpolar barrier for these viruses and revealed size fraction-dependent Arctic diversity hotspots for some virus groups, which may reflect a high diversity of their hosts in cold environments. Temperature was an important factor that shaped the virus–host interactions of polar environments. Consistent with these findings, our analyses suggested a presently unidentified virus–host relationship between polar diatoms and giant viruses. Our phylogenomic tree and ancestral state reconstruction revealed back-and-forth adaptations between lower- and higher-temperature niches that occurred recurrently throughout the long evolutionary course of these viruses. Numerous functions, especially ones related to host interactions, were found to be specific to viral polar adaptation, but most of them were not identified as polar-specific functions in eukaryotes. Furthermore, the gene repertoire of these large DNA viral genomes appears more evolutionarily flexible and responsive to temperature change than that of eukaryotic genomes. The discovery of this difference in gene repertoire between polar and nonpolar viruses infecting marine eukaryotes prompts concern about the influence of climate warming on the marine ecosystem, given the importance of these viruses in regulating their host communities and biogeochemical cycling.

## Methods

### Global Ocean Eukaryotic Viral (GOEV) database

Metagenomic datasets and environmental data are provided in Supplementary Table 1. The Global Ocean Eukaryotic Viral (GOEV) database contained 1,817 viral genomes^28,30,63^. Taxonomic inference, read mapping, gene call and gene annotation of the GOEV were performed in a previous work^28^. 1380 detected viruses were classified into six main taxonomic groups: five orders (i.e., *Algavirales, Asfuvirales, Imitervirales, Pandoravirales*, and *Pimascovirales*) and the newly discovered phylum, *Mirusviricota*. Six different size fractions were used in this study: 0.22–1.6 μm or 0.22–3.0 μm (“Pico”), 0.8–5 μm (“Piconano”), 5–20 μm (“Nano”), 20–200 μm (“Micro”), 200–2,000 μm (“Macro”), and 0.8–2,000 μm (“Broad”). The size fraction below 0.22 μm was excluded in this study because of the low relative abundance and high overlap with species from the Pico size fraction. Mean coverage of these viruses was transformed into RPKM (Reads Per Kilobase of exon per Million mapped reads) using the formula: numReads / (genomeLength/1000 * totalNumReads/1,000,000). RPKM profile was used for the ecological analyses in this study.

### Phylogenetic tree construction

Phylogenetic trees used in this study were reconstructed using IQ-TREE v.1.6.2^64^. The viral species tree was reconstructed with the site-specific frequency PMSF model following a best-fitting model according to the BIC from the ModelFinder Plus option. The PolB tree was of *Nucleocytoviricota* reference genomes and *Chaetoceros* genomes was reconstructed with the LG+F+I+G4 model. Tree structure manipulation and analysis were done using ETE3 toolkit v.3.1.1^65^. iTOL v.6 was used to visualize the phylogenetic trees^66^. Phylogenetic diversity was calculated using the ‘pd’ function in the R package ‘picante’^67^ for polar and nonpolar genome subsets.

### Ecological analyses

Diversity analyses were performed using R v.4.0.1^68^ in Rstudio v.1.3.959^69^. To evaluate the diversity of each sample, the richness (number of MAGs), Shannon’s index and Pielou’s evenness were calculated with the package ‘`vegan`’^70^. Compositional variation among samples was assessed with a non-metric multidimensional scaling (NMDS) ordination based on Bray-Curtis dissimilarity. Samples with low viral abundance and richness produce outliers that reduce the readability of the NMDS ordination plot. To avoid such a bias, samples for which the sum of cumulative coverage was less than 10 or richness was less than 5 (set as the cutoff threshold) were removed from the compositional variation analyses. Statistical significance of differences among the sample groups (size fractions and biomes) was tested using an ANOSIM (analysis of similarities) with 9,999 permutations. The significance threshold was set to a *p*-value of 0.01. The plots and maps of sampling stations were generated by packages ‘**`ggplot2`**’^71^ and ‘**`rgdal`**’^72^.

### Gene annotation and clustering

Genes were predicted using Prodigal v.2.6.3^73^ within anvi’o v6.1^74^ with the default parameters. Gene cluster communities were classified through the AGNOSTOS^75^ workflow. Those two steps were performed and described in a previous work^28^. For functional annotation, genes were assigned to KEGG Orthologs (KOs) using eggNOG-mapper v.2.1.5^76^ (“Diamond” with an E-value cut-off of 1.0×10^−5^). Viral marker genes were searched with in-house HMM profiles from NCVOG (nucleocytoplasmic virus orthologous genes)^77^ and GVOG (giant virus orthologous groups)^22^ databases using HMMER v.3.2.1 (http://hmmer.org) with an E-value of 1×10^−3^. ^78^Antifreeze proteins were detected using InterProScan v.5.44-79.0^78^. Low-complexity regions of protein sequences were identified using the option ‘-qmask seg’ in usearch v.11.0.667^79^.

### Virus–plankton interaction network

We determined the relative abundance matrix for the virus MAGs from the Pico size fractions and relative abundance matrices for eukaryotic MAGs from five cellular size fractions (Piconano, Nano, Micro, Macro, and Broad). To create the input files for network inference, we combined the viral matrix with each of the eukaryotic matrices (corresponding to different size fractions), and only the samples represented by both viral and eukaryotic MAGs were placed in new files. Relative abundances in the newly-generated matrices were normalized using centred log-ratio (*clr*) transformation after adding a pseudo-count of one to all matrix elements because zero cannot be transformed in *clr*. Normalization and filtering were separately applied to viral and eukaryotic MAGs.We then removed the MAGs that had fewer than three sample observations. Network inference was performed using FlashWeave v.0.15.0^80^ with Sensitive mode to set a threshold of α < 0.01 as the statistical significance and without the default normalization step. All detected pairwise associations were then assigned a weight that ranged between −1 and +1. The network was visualized with Cytoscape v.3.7.1^81^ using the prefuse force-directed layout. Proteins between linked genome pairs were aligned using BlastP in Diamond v.2.0.6^82^ with an E-value cut-off of 1.0×10^−50^.

### Host prediction

First, we pooled network associations from five size fractions by keeping the best positive or negative associations (i.e., the edges with the highest absolute weights). We used a phylogeny-guided filtering approach, Taxon Interaction Mapper (TIM)^37^, to predict the host using the global nucleocytoplasmic large DNA virus (NCLDV)–eukaryote network. TIM provides a list of nodes in the viral tree and associated NCBI taxonomies (order, class, and phylum) of eukaryotes that show significant enrichment in the leaves under the nodes. All the virus–eukaryote associations were mapped on the viral phylogenetic tree to calculate the significance of the enrichment of specific associations using TIM, and the result was visualized with iTOL v.6.

### Endogenous viral signals

We searched the viral signals in 713 genomes from the eukaryotic environmental genomes database using VirSorter2 v.2.2.3^29,83^. Both NCLDV and *Lavidaviridae* (virophage) genomic insertions (or co-binning) were searched using --min-score 0.85 and 0.95 for NCLDV and virophage, respectively. We next obtained long-read assembled genomes of two *Chaetoceros* isolates, *C. muelleri* ^52^and *C. tenuissimus* ^51^. Giant Endogenous Viral Elements (GEVEs) were detected using ViralRecall v.2.1 (-s 5 -w 10)^84^. *Nucleocytoviricota* DNApolB sequences in *Chaetoceros* genomes were detected using HMMER v.3.2.1 search against an in-house DNApolB database. *Chaetoceros*-originating DNApolB sequences were manually concatenated if they were in the same contig and had continuous gene IDs. *Chaetoceros*-originating DNApolBs were aligned with other reference NCLDV DNApolBs using MAFFT-linsi v.7.453^85^, and the phylogenetic tree was constructed using IQ-TREE as described above^64^.

### Size index

Each *Tara* Oceans metagenome corresponds to a specific filtering size fraction (Pico, Piconano, Nano, Micro, Macro, and Broad size fractions as defined above), which were sorted as a list by increasing size. An index constant was set for each size fraction from small to large: Pico = 1, Piconano & Broad = 2, Nano = 3, Micro = 4, Macro = 5 (the Broad and Piconano size fractions were merged because of their similar relative abundances and lack of Arctic samples for the Piconano fraction). We calculated the size index for a given genome by first multiplying the RPKM of the genome in a sample by the corresponding index constant, then dividing the sum of the products by the overall sum of the RPKMs of the genomes from all samples.

### Biome and size niche

Each sample was associated with one specific marine biome (Coastal, Trades, Westerlies, or Polar). To investigate the difference between polar and nonpolar regions, we pooled Coastal, Trades, and Westerlies samples as “Nonpolar”. First, we assigned each genome to Polar or Nonpolar if a genome was exclusive to either nonpolar or polar biomes. Additionally, on the basis of RPKM profiles, we calculated the significance using the Wilcoxon rank-sum test. Adjustments for multiple testing were performed using the Benjamini-Hochberg (BH). The significance threshold was set to a corrected *P*-value of 0.05. Similar assignments were performed for two size fractions: intercellular (Pico-size) and intracellular (Piconano, Nano, Micro, Macro, and Broad).

### Robust ecological optimum and tolerance

We calculated the robust ecological optimum for a genome (or a gene), which reflects the optimal living condition regarding a given environmental parameter and a tolerance range around this optimum defined by lower and upper bounds^32,33^. For each genome (or a gene), we computed the proportion of RPKM in a given sample relative to the sum of RPKM over all samples. We then used these proportions to populate a weighted vector of a fixed size (n = 10,000) with environmental values accordingly. The ecological optimum is then defined as the median value (Q2) of this vector, and the tolerance (niche) range is given by the interquartile range (Q3 to Q1; some environmental parameter values were missing [nonavailable (NA)] for some samples). To avoid inferring spurious ecological optima and tolerance ranges for genomes (or genes) for which there were many missing values, we set a minimum threshold of 10 observations with non-NAs and a minimum fraction of 30% non-NA values.

### Ancestral states estimation and Relative Evolution Divergency

Ancestral states of Nonpolar and Polar viruses were estimated using the function “`ace`” (Ancestral Character Estimation) in the R package ‘`ape`’^86^. The input files were a rooted phylogenetic tree based on the four-hallmark gene set described above. In the tree, we retained only viruses with biome assignments of Polar or Nonpolar, and excluded viruses with “Unknown” biomes. We used `type = “discrete”, method = “ML”`, and `model = “ER”` (one-parameter equal rates model). The ancestral states were analysed based on a series of likelihood values for Polar and Nonpolar. Relative Evolutionary Divergence (RED) values were calculated using the “`get_reds`” function in the package “`castor`”^87^.

### KO enrichment in Polar viral genomes

“Polar”, “Nonpolar”, or “Unknown” biome niche was assigned to each viral genome as described previously. For individual lineages at four taxonomic levels (root, main group, family, and genus), the enrichment of a given KO in Polar genomes assessed using Fisher’s exact test in SciPy v.1.7.1^88^. Adjustments for multiple testing were performed using the Benjamini-Hochberg (BH). The significance threshold was set to a corrected *P*-value of 0.05.

### Phylogenetic signal of functions

We hypothesized that the phylogenetic distributions of some polar specific functions (i.e., “trait distribution”) could be better explained in part by environment selection rather than only by speciation history. We therefore compared two models, (i) the Brownian motion model (Pagel’s lambda = 1, used as the null hypothesis in which the distribution of a trait is simply explained by species tree) and (ii) the Lambda model (0 ≤ Pagel’s lambda ≤ 1; lambda = 0 corresponds to the lack of phylogenetic signal in the distribution of a trait), by the likelihood ratio test using the function “`fitContinuous`” in an R package “`geiger`”^89^. The *p*-values to reject the null hypothesis were calculated by assuming chi-squared distribution with 1 d.f. for the likelihood-ratio test statistic and adjusted using the BH procedure. The threshold was set to a corrected *p*-value of 0.05

## Acknowledgements

This work was supported by JSPS/KAKENHI (18H02279 and 22H00384, to H. O.), and the Collaborative International Joint Research Program of the Institute for Chemical Research, Kyoto University (No. 2021-29, 2022-26 to T.O.D.; No. 2022-27, to S. C.), and the H2020 European Commission project AtlantECO (award number 862923, to S. C.). Computational time was provided by the SuperComputer System, Institute for Chemical Research, Kyoto University. We further thank the *Tara* Oceans consortium, and the people and sponsors who supported *Tara* Oceans. *Tara* Oceans (including both the *Tara* Oceans and *Tara* Oceans Polar Circle expeditions) would not exist without the leadership of the *Tara* Expeditions Foundation and the continuous support of 23 institutes (https://oceans.taraexpeditions.org). This article is contribution number XXX of *Tara* Oceans. We thank Gabe Yedid, PhD, from Edanz (http://jp.edanz.com) for editing a draft of this manuscript.

## Author contributions

L. M. and H. O. designed the study. L. M. performed the primary biogeographical analysis. T.O.D completed the genome-resolved metagenomic analysis. M. G. performed phylogenomic analyses. E. P. generated the reads mapping data. A. F-G provided de novo clusters of viral genes. R.Y.N, J. W, H. K. contributed to the bioinformatics analysis. All the authors contributed to interpreting the data and writing the manuscript.

