## Supplementary_Text for "Genomic adaptation of giant viruses in polar oceans"

**Biogeography of giant virus genomes**

We used giant virus genomes from the Global Ocean Eukaryotic Viral (GOEV) database[1]. Their abundance profiles across *Tara* Oceans samples from different size fractions (Supplementary Fig. 1a,b; Supplementary Table 1) revealed 1,380 viral genomes that showed signals in at least one sample out of 928 samples (see Methods; Supplementary Fig. 1a,b). Rarefaction analyses showed reached a plateau when all *Tara* Oceans samples are combined, but genomes in micro- (20-200 μm) and macro-size (200-2000 μm) fractions were still under-sampled (Supplementary Fig. 2a,b). Detected viruses were taxonomically classified into six main groups based on our phylogenomic analysis, with genome sizes ranging from 50 Kb to 1.6 Mb: *Algavirales* (n = 155), *Asfuvirales* (n = 9), *Imitervirales* (n = 913), *Pandoravirales* (n = 81), *Pimascovirales* (n = 75), and *Mirusviricota* (n = 111). Of these, all but the *Mirusviricota* belong to the *Nucleocytoviricota* phylum of the realm *Varidnaviria*. *Mirusviricota* belongs to the realm *Duplodnaviria* (Supplementary Table 2). These viruses, recently identified, possess indeed the morphogenetic module of this realm, even though a much larger fraction of informational genes relates them to *Nucleocytoviricota*[1].

In agreement with previous results based on marker genes[2,3], imiterviruses formed the largest group, with some members distributed widely (>100 samples) but at relatively low abundance (Supplementary Fig. 1c; Supplementary Fig. 2c,d). The cumulative coverage (i.e., sum of the coverages in all samples for a given genome) in the second largest group, algaviruses, was significantly higher than for other groups (Kruskal-Wallis test, $P = 1.47\times10^{-21}$) but these viruses were observed in fewer samples (≤95 samples) than *Imitervirales* (Supplementary Fig. 2c; Supplementary Fig. 2d).

Most previous metagenomic surveys on giant viruses analysed data from a pico-size fraction (0.2–3.0 μm)[4,5]. Our dataset allowed us to assess abundance of viruses in larger size fractions up to the meso-fraction (200–2000 μm). Giant viruses were detected in both small and large size fractions at many stations (Supplementary Fig. 1b). Accordingly, size index values (i.e., a measure of sampling size fraction preference for each genome) were distributed widely for giant viruses compared with those of individual eukaryotic taxa (Supplementary Fig. 3a,b). Viral signals in large size fractions (e.g., >0.8 μm) may originate from viral genomes inside their host cells, whereas those in small size fractions (e.g., 0.2–3.0 μm) may originate from viral genomes from either free virions or within host cells.

We assigned infection stage categories to individual viral genomes [either "virion" (0.2–3.0 μm) or "cellular" (>0.8 μm)] based on the reads per kilobase of genome per million reads mapped (RPKM) distribution across size fractions (Supplementary Fig. 3c; Supplementary Table 2). This categorization demonstrated that 15% of viruses (n = 211) were over-represented in the "virion" category, while 8% (n = 111) were over-represented in the "cellular" category. The proportions of *Imitervirales* and *Algavirales* were relatively low, and those of *Pimascovirales*, *Pandoravirales*, *Asfuvirales* and *Mirusviricota* were relatively high for the "cellular" category compared with those in the "virion" category, implying different host size ranges for different groups of viruses.
