## Supplementery_Figures for "Genomic adaptation of giant viruses in polar oceans"

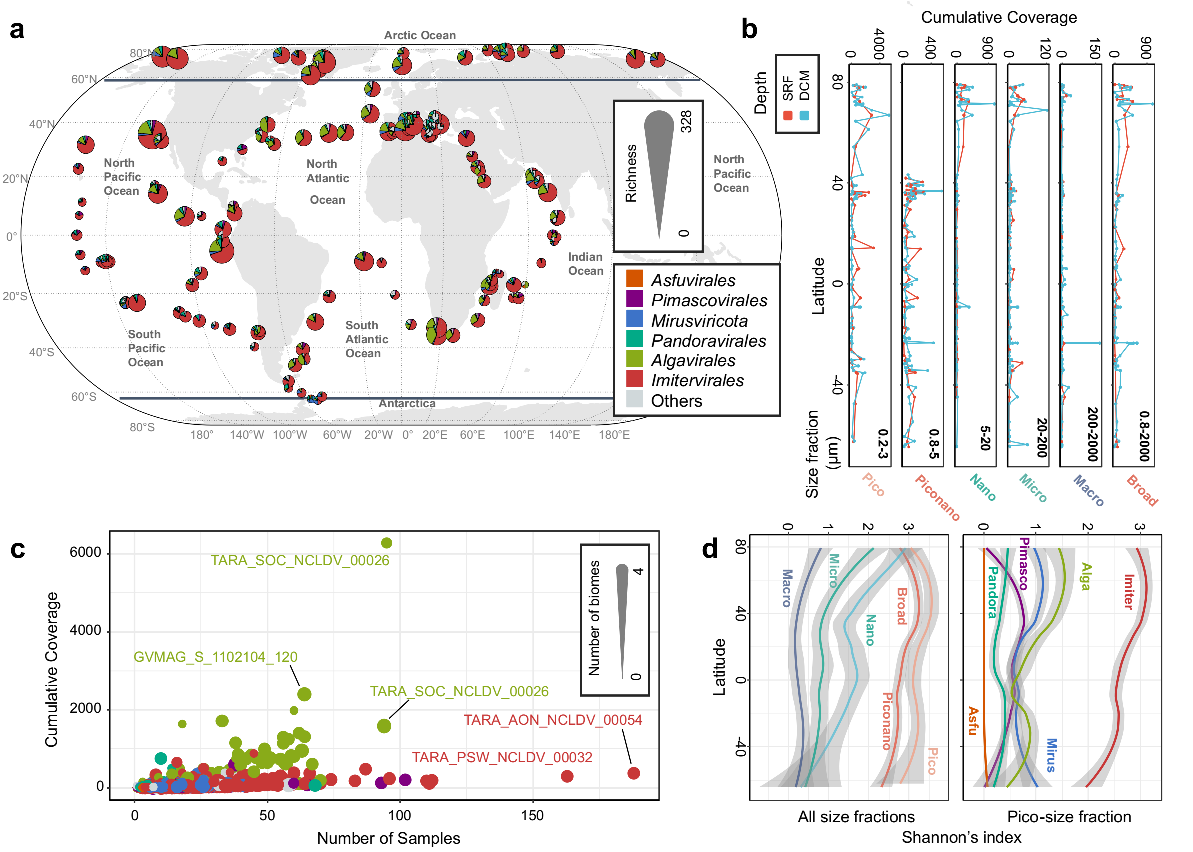


**Supplementary Fig. 1 | Distribution of giant virus genomes. a,** Richness of viral communities at the stations of the *Tara* Oceans expedition (2009–2013). A total of 928 epipelagic metagenomes from 143 *Tara* Oceans stations are included in this study. Each pie chart represents the contributions to richness by six taxonomic main groups, and the size of the pie chart is proportional to the total richness at the station. The richness of two depths (Surface and Deep Chlorophyll Maximum) and different size fractions (Pico, Piconano, Nano, Micro, Macro, and Broad) are integrated into one pie chart. Dashed lines indicate the boundary of Polar samples. **b,** Cumulative coverage in each sample along the latitude. Individual size fractions are displayed separately, and line colours correspond to two depths (red for Surface, and blue for Deep Chlorophyll Maximum). **c,** Cumulative coverage of individual viral genomes versus the number of samples in which the respective genomes were observed. The top five abundant/prevalent genomes are labelled. **d,** Locally estimated scatterplot smoothing plots of the latitudinal distributions of viral diversity (Shannon's index). The left panel shows the pattern of diversity in each size fraction. The Broad and Piconano size fractions were pooled because of their similar relative abundances and lack of Arctic samples in the Piconano size fraction. The right panel shows the diversity of communities of six main groups in the Pico, Piconano, and Broad size fractions.

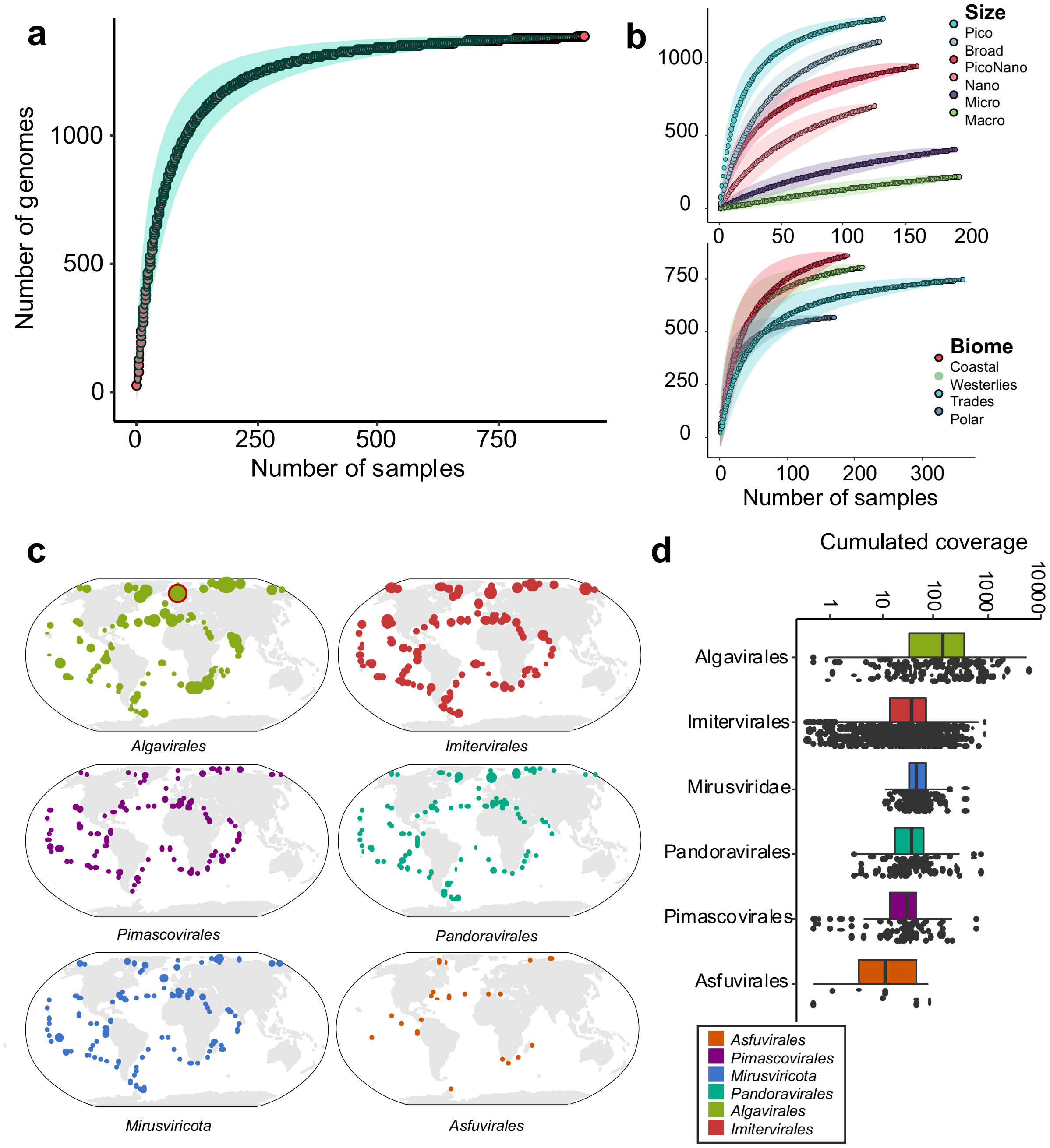


**Supplementary Fig. 2 | Summary of the biogeography of viruses. a,** Rarefaction curves for marine eukaryotic viral genomes in all samples and **b,** in subsamples separated by size fractions and biomes. For each curve, the average of 100 permutations is displayed with dots and the standard deviation is displayed with colour ranges. Rarefaction analyses reached a plateau when all *Tara* Oceans samples were combined, but genomes in Micro (20–200 µm) and Macro (200–2000 µm) size fractions were still undersampled. **c,** Maps of the cumulative coverage of giant viruses in six main groups. Dot sizes are normalized by that of the most abundant dot, *Algavirales* in station 188 (marked with a red framed circle). **d,** Jitter box plots for the cumulative coverage of viral genomes classified into six main groups. There was a significant difference between groups ($P = 1.47 \times 10^{-21}$, Kruskal-Wallis test).

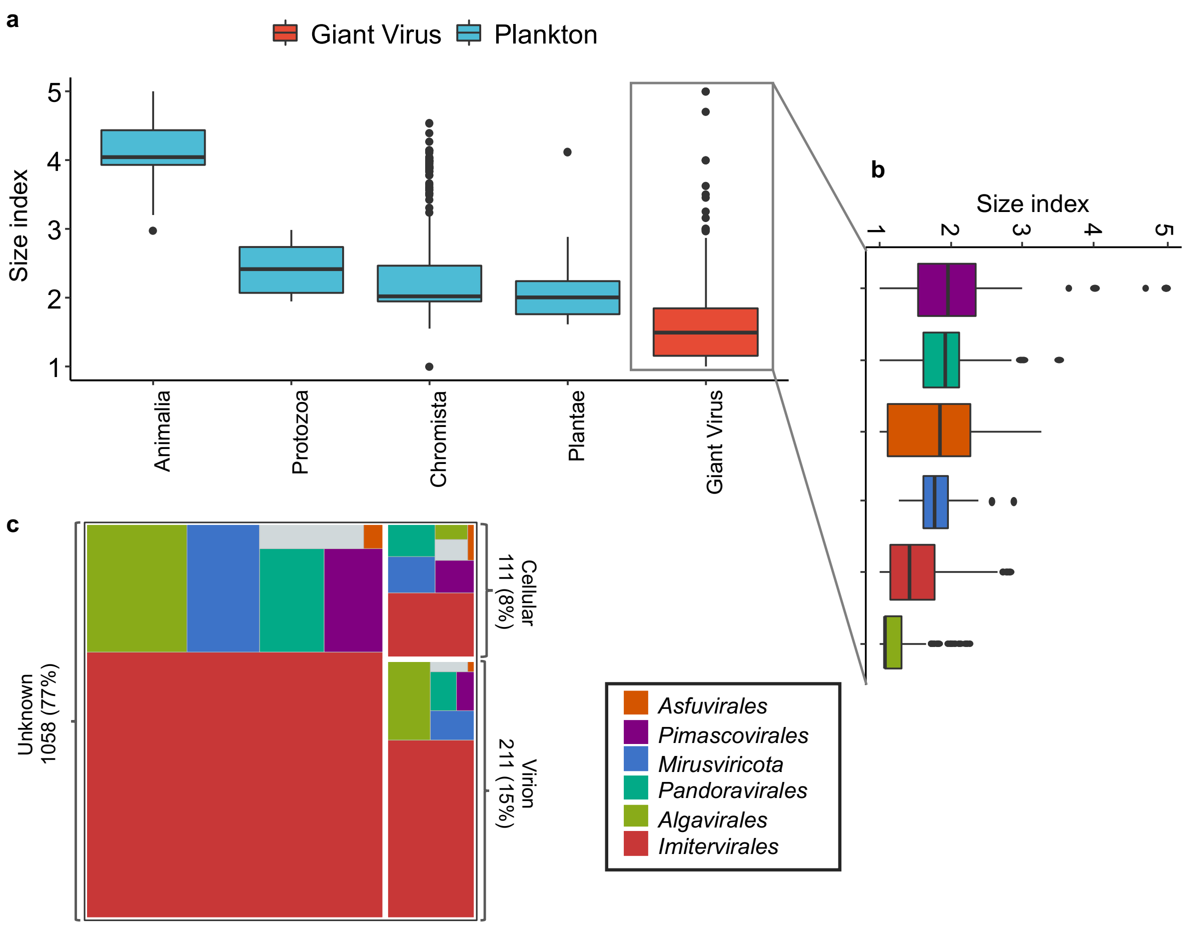


**Supplementary Fig. 3 | Size distribution of viruses. a,** Boxplots for the size indices of eukaryotic kingdoms and giant viruses. **b,** Boxplots for the size indices of six virus main groups. **c,** Treemap diagram showing the number of giant viruses assigned to "virion" or "cellular" size categories. Colours indicate the main groups.

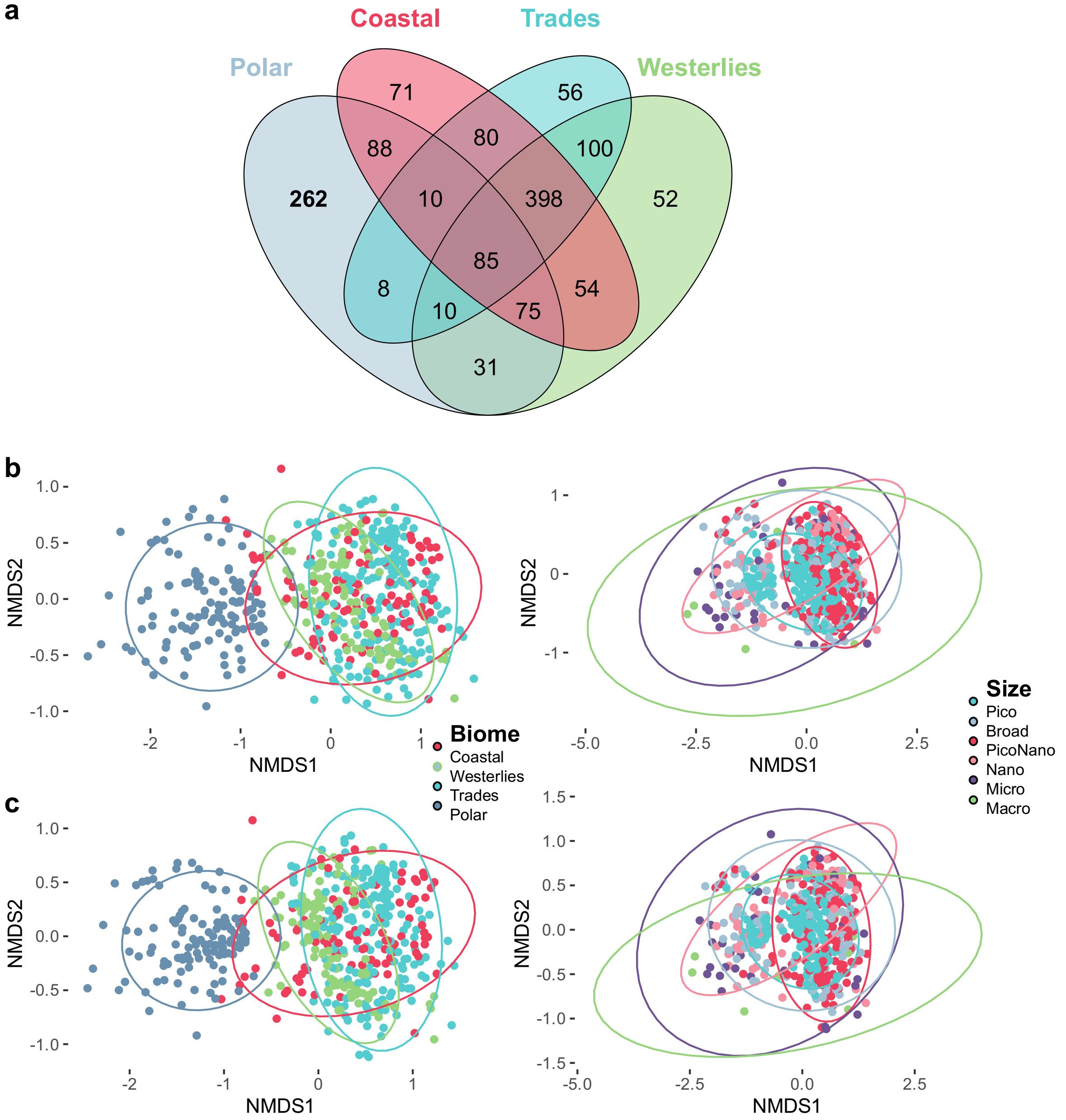


**Supplementary Fig. 4 | Community structures of viruses. a,** Venn diagram showing the numbers of shared or unique giant viruses across biomes. **b,c** NMDS ordination (Bray–Curtis dissimilarity) of the viruses communities for all samples using **(b)** relative abundance data (stress = 0.1617) and **(c)** presence/absence data (stress = 0.1541). Colours indicate biomes (left) and size fractions (right). Ellipses represent 95% confidence levels for each group. Statistical significance of the groupings was confirmed for both biome and size fraction groups (ANOSIM, $P < 0.01$)

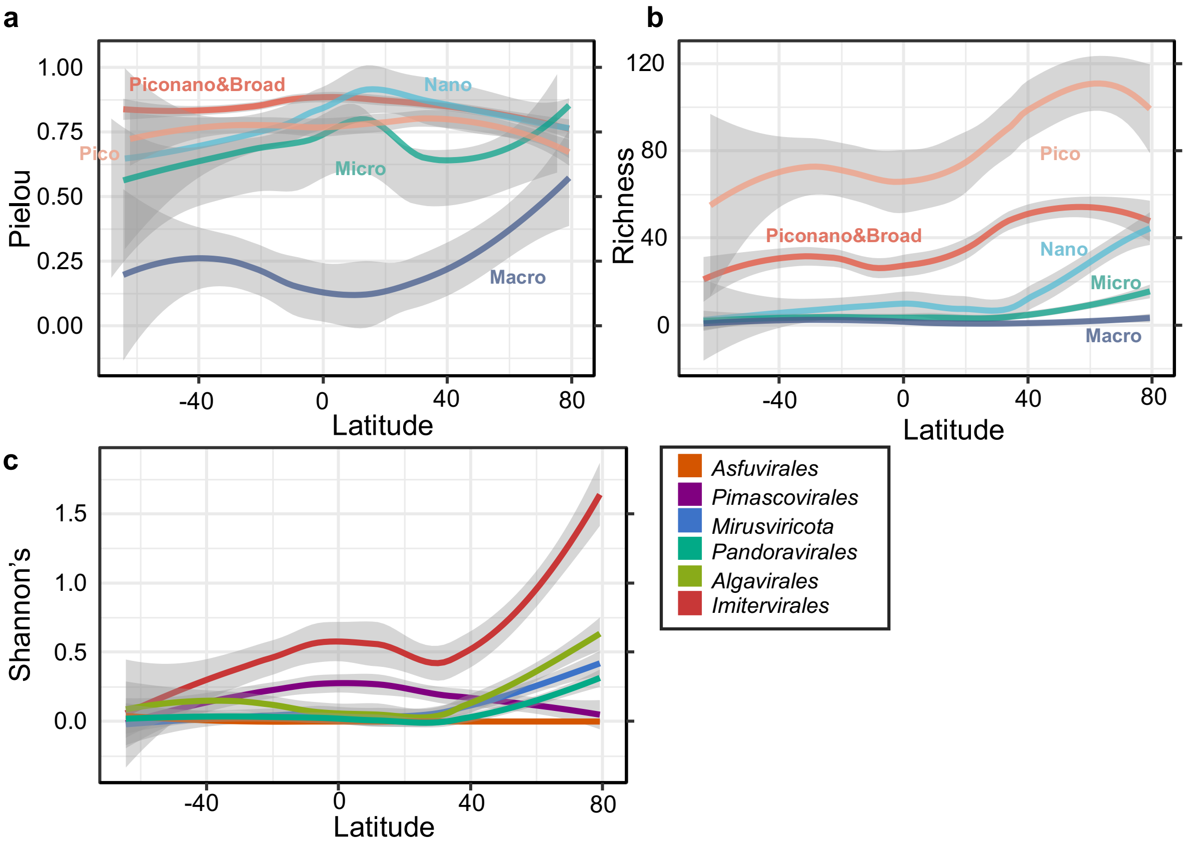


**Supplementary Fig. 5 | Locally estimated scatterplot smoothing plots of the latitudinal distributions of viral communities.** Latitudinal variation in **a,** Pielou's evenness and **b,** richness of marine viral communities for different size fractions. Colours indicate size fractions. **c,** Shannon's index of communities of six main groups in large-size fractions (Nano, Micro, and Macro size fractions). Shaded areas represent 95% confidence intervals.

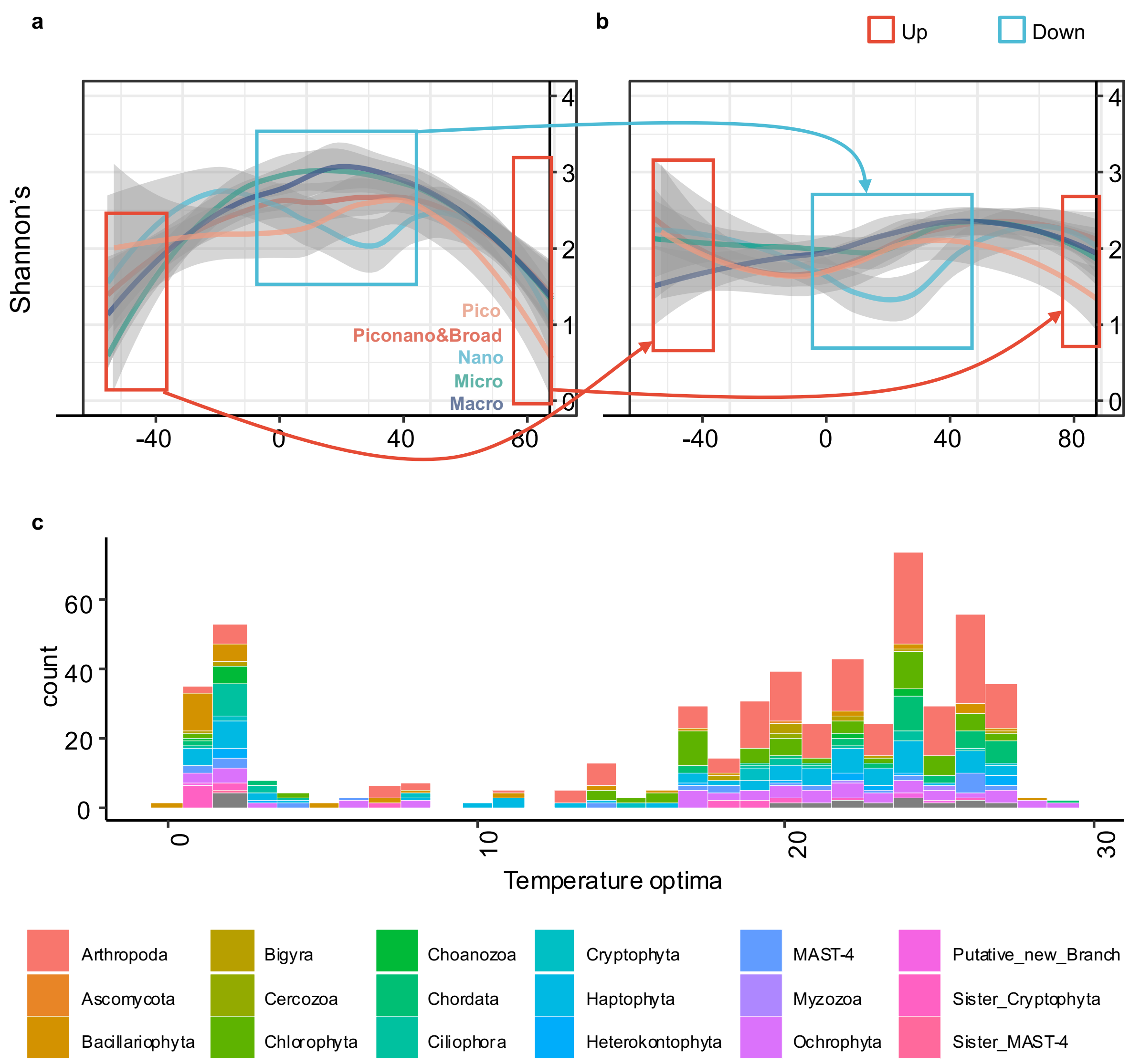


**Supplementary Fig. 6 | Eukaryotic communities in the virus-host interactome**. **a,b** Locally estimated scatterplot smoothing plots of the latitudinal distributions of eukaryotic community diversity (Shannon's index). **a,** Diversity of communities of eukaryotes that have no association with viruses in the network. **b,** Diversity of communities of eukaryotes that have associations with viruses in the network. Colours stand for size fractions. Polar regions and equatorial regions were marked with coloured frames. (C) Histogram of temperature optima of eukaryotic nodes. Colours represent the phylum of eukaryotic nodes as indicated at the bottom of the plots. Eukaryotic nodes with low temperature optima were enriched in diatoms (Bacillariophyta).

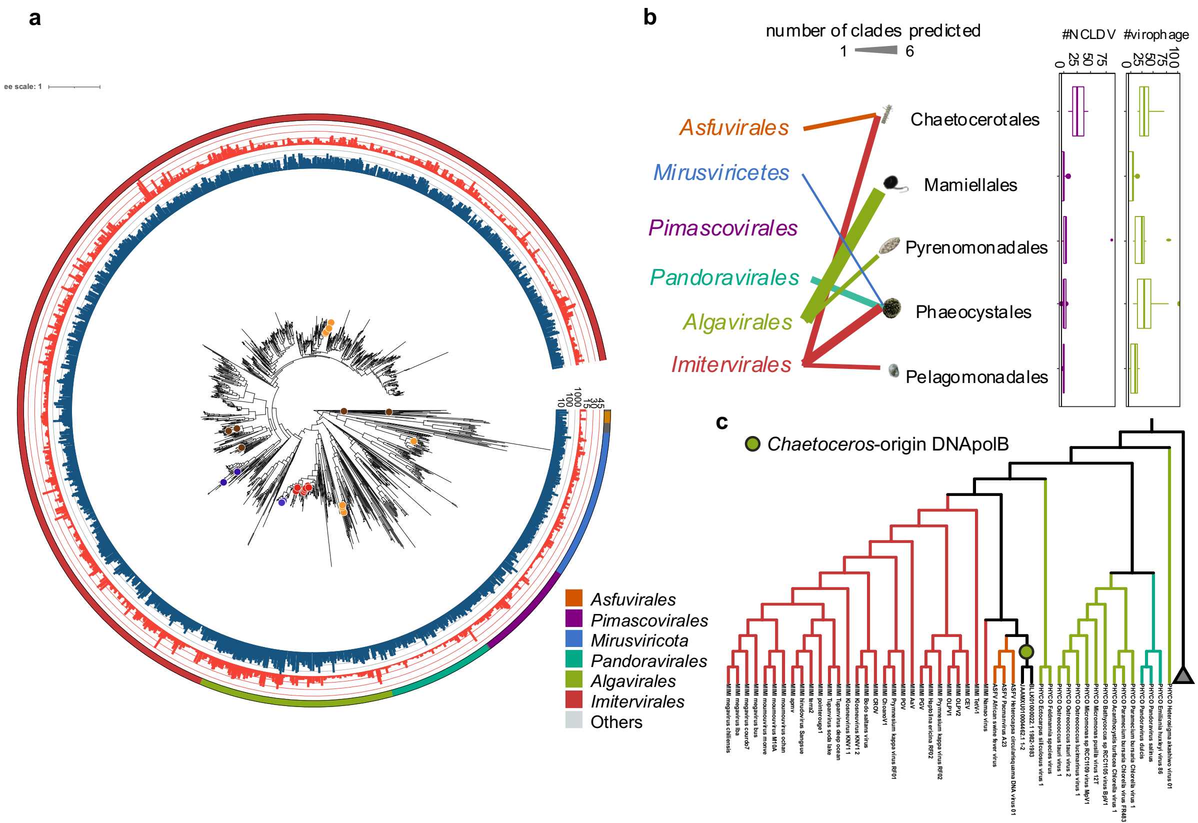


**Supplementary Fig. 7 | Host prediction of viruses. a,** Phylogenetic tree based on four hallmark genes with putative host groups (coloured circles) predicted by TIM, which are summarized in (**b**). The outermost layer shows the taxonomy of six main groups. The middle and inner layers show the number of stations in which the viruses were observed and cumulative coverage, respectively. **b,** Summary of host prediction results. Left panel: line colours represent the six main groups; line widths are proportional to the number of clades predicted to the associated hosts. Right panel: boxplots show the number of viral insertions detected in predicted host genomes. **c,** Phylogenetic tree constructed from PolB sequences from the *Nucleocytoviricota* reference genomes and PolB homologs from two *Chaetoceros* isolate genomes. Viral gene sequences are coloured according to the main group classification, and the sequences of *Chaetoceros* are marked using a green circle. The superclade "MAPI" was collapsed and used as the outgroup.

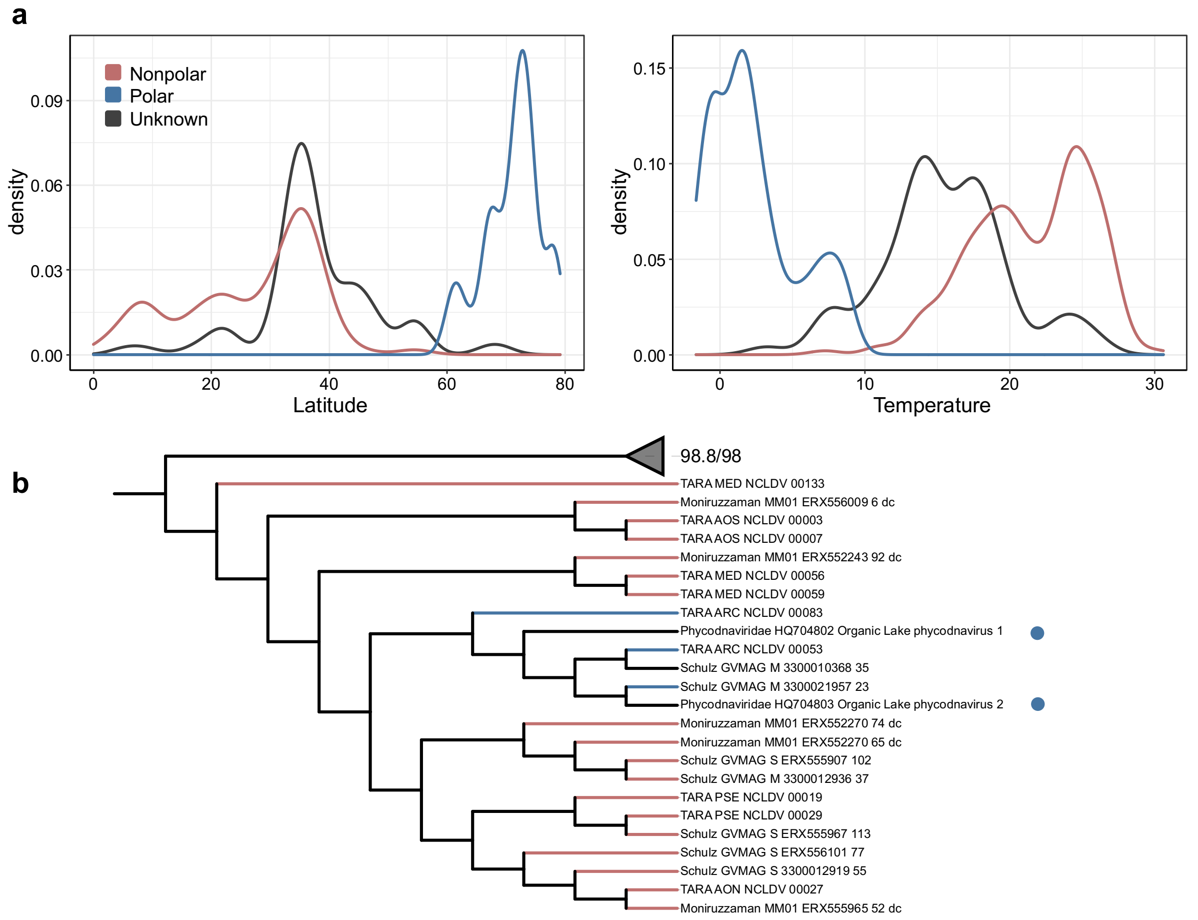


**Supplementary Fig. 8 | Niche assignments of viral genomes. a,** Density plots of the latitude and temperature optima. Three biome assignment groups (Polar, Nonpolar, and Unknown) are displayed separately. **b,** The Polar clade containing the reference Organic Lake phycodnaviruses (blue circles). Ancestral states of Nonpolar and Polar were estimated using the phylogenetic tree based on a one-parameter equal rates model. Blue stands for Polar and red stands for Nonpolar viruses.

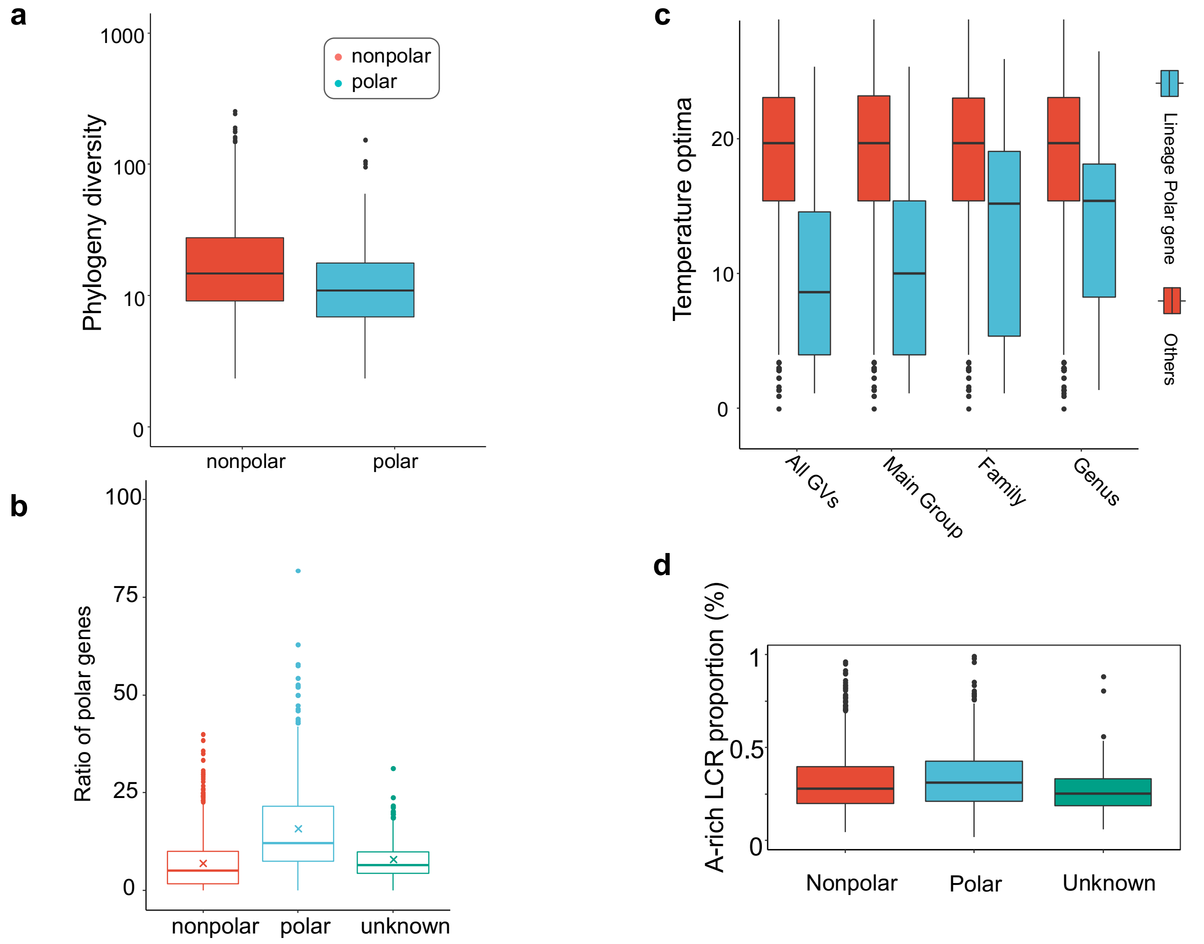


**Supplementary Fig. 9 | Polar-specific KOs in polar viral genomes. a,** Phylogenetic diversity of Polar-specific and Nonpolar-specific KOs. **b,** Ratio of Polar-specific KOs in Polar, Nonpolar and biome-unknown genomes. **c,** Boxplots of the temperature optima of Polar KOs enriched in Polar viral genomes at least one lineage (blue) and other KOs (KOs that were not enriched in Polar viral genomes at any lineage at four taxonomy levels, i.e., root, main group, family, and genus) (red). Enrichment analyses were performed at the main group, family, and genus levels. **d,** Alanine(A)-rich low-complexity regions (LCR) proportion in Polar, Nonpolar and biome-unknown genomes.

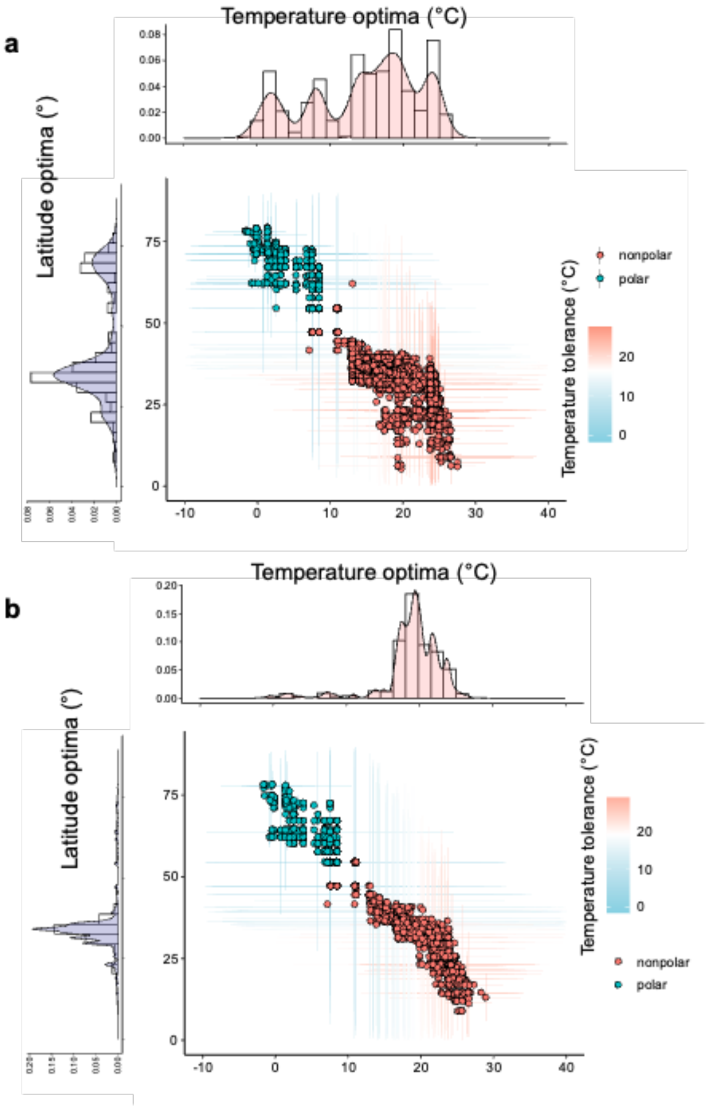


**Supplementary Fig. 10 | Distributions of robust ecological optima for the orthologous groups of viruses and KOs in eukaryotic genomes. a,** Scatterplot of the temperature and latitude optimum of AGNOSTOS gene cluster communities (GCCs) for viruses. Bars indicate the tolerance ranges of temperature (horizontal) and latitude (vertical). Histograms show the distributions of the temperature and latitude optima of GCCs. **b,** Distribution of temperature and latitude optima of KOs of eukaryotic genomes.
